## Supplementary figures and images for "Optogenetic Miro cleavage reveals direct consequences of real-time loss of function in *Drosophila*"

### Supporting Figure 1

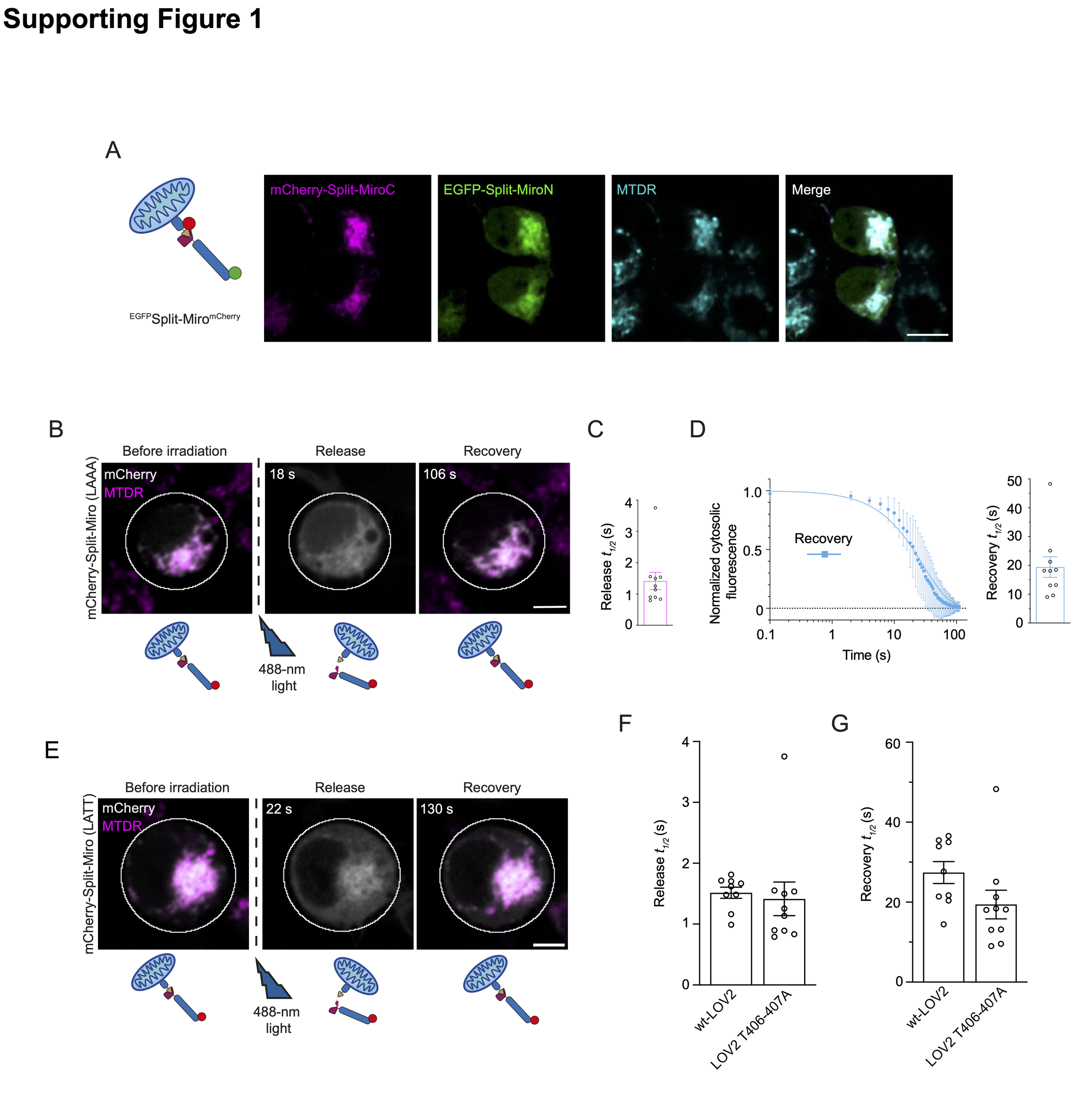

### Supporting Figure 2

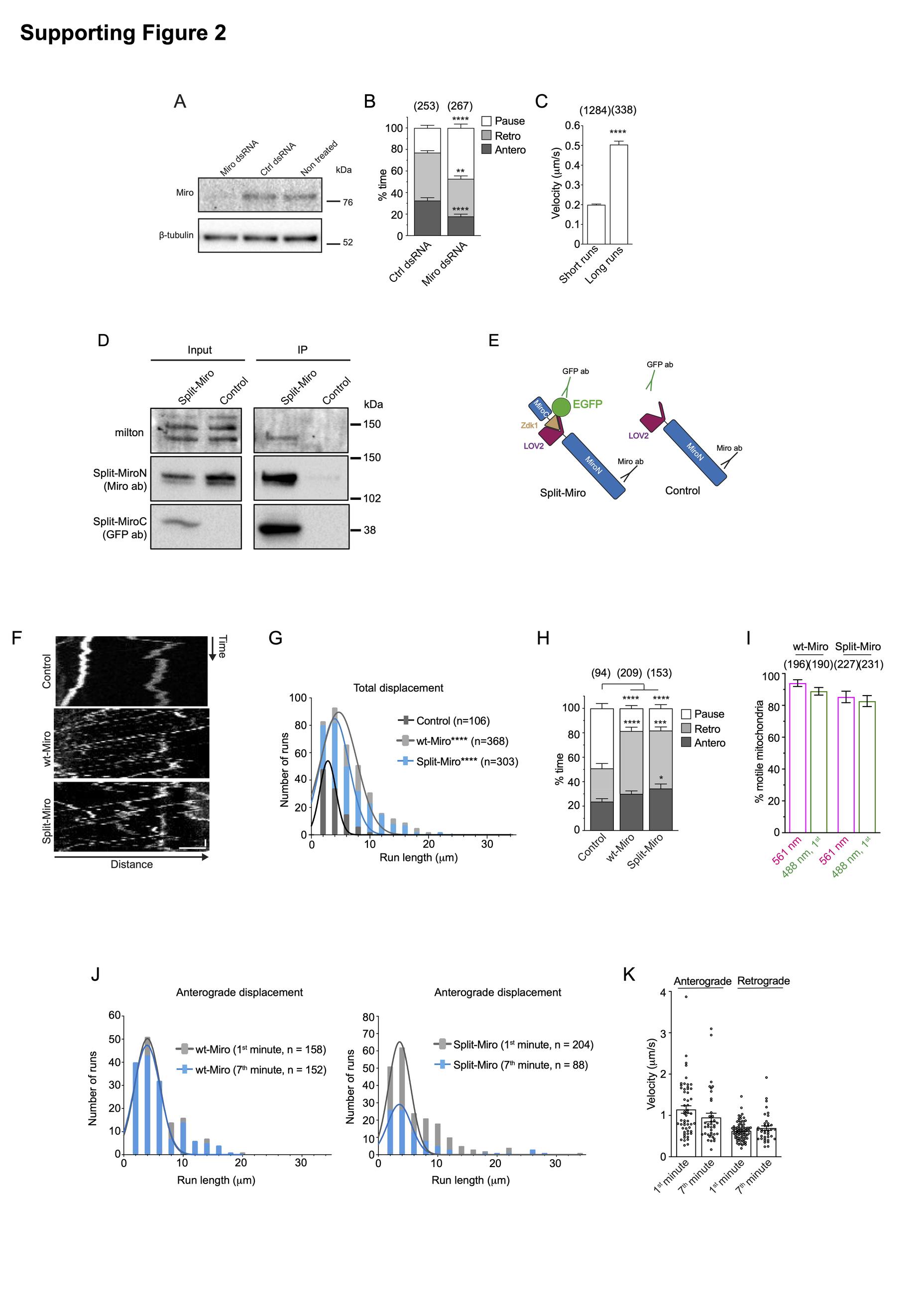

### Supporting Figure 3

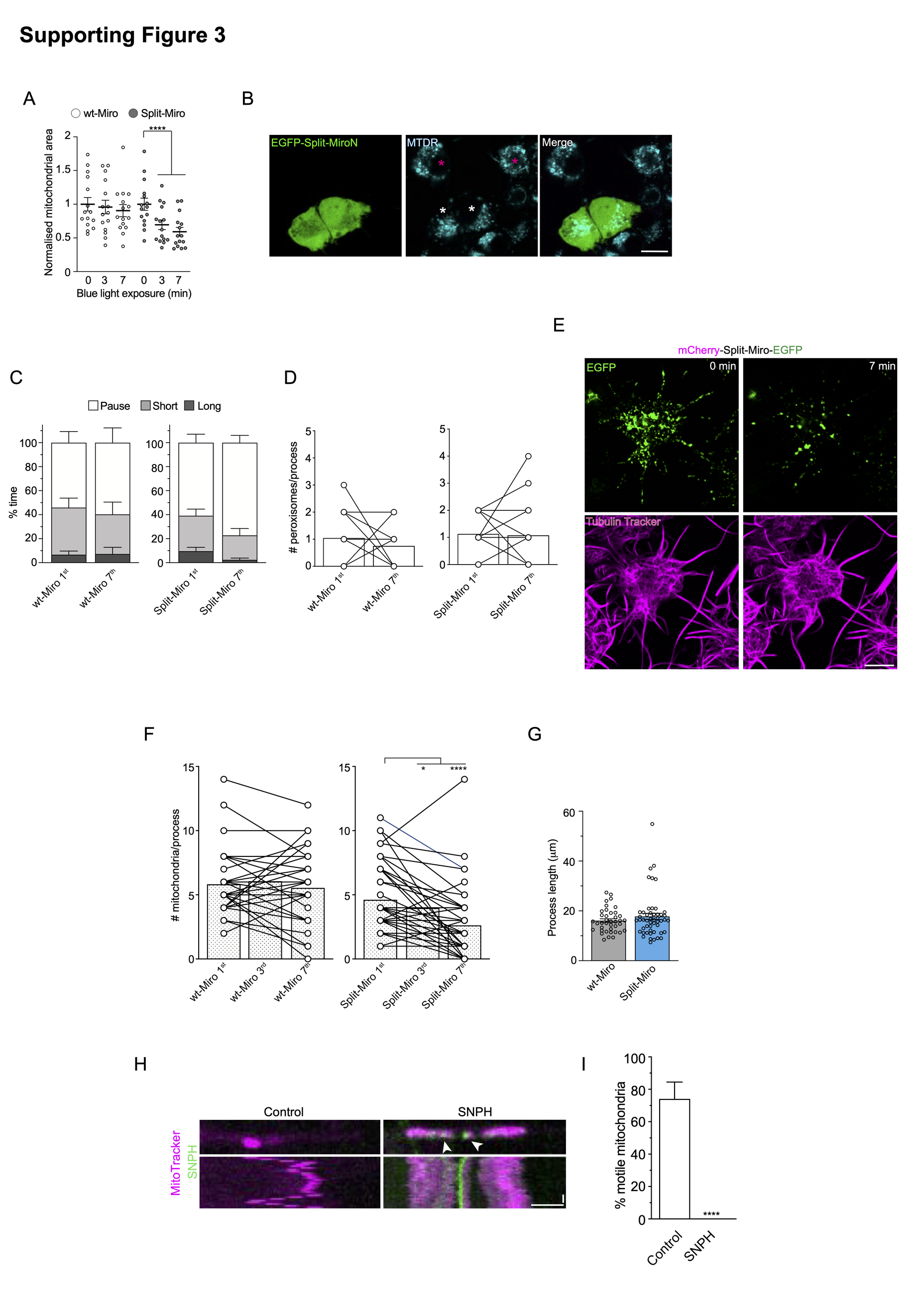

### Supporting Figure 4

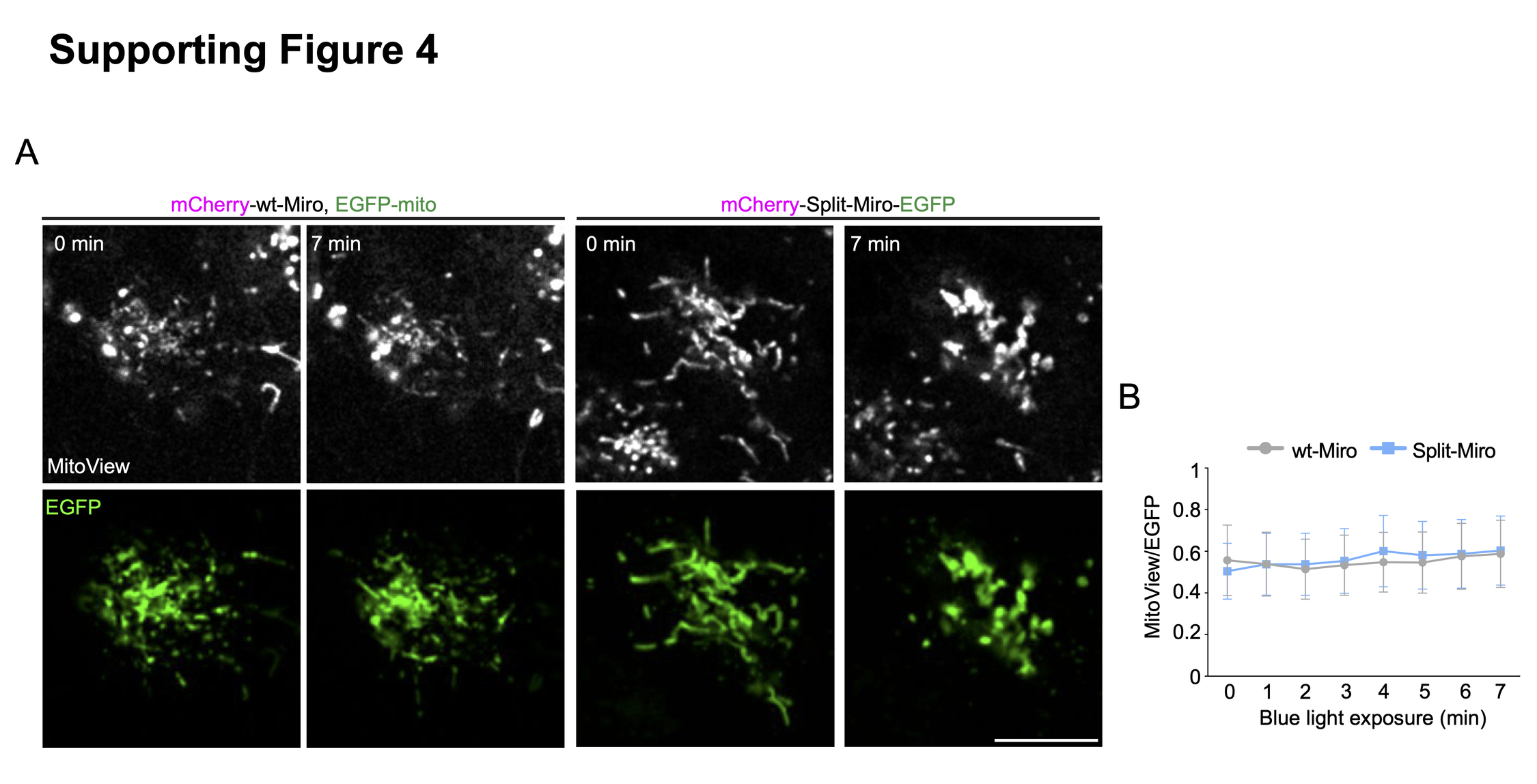

### Supporting Figure 5

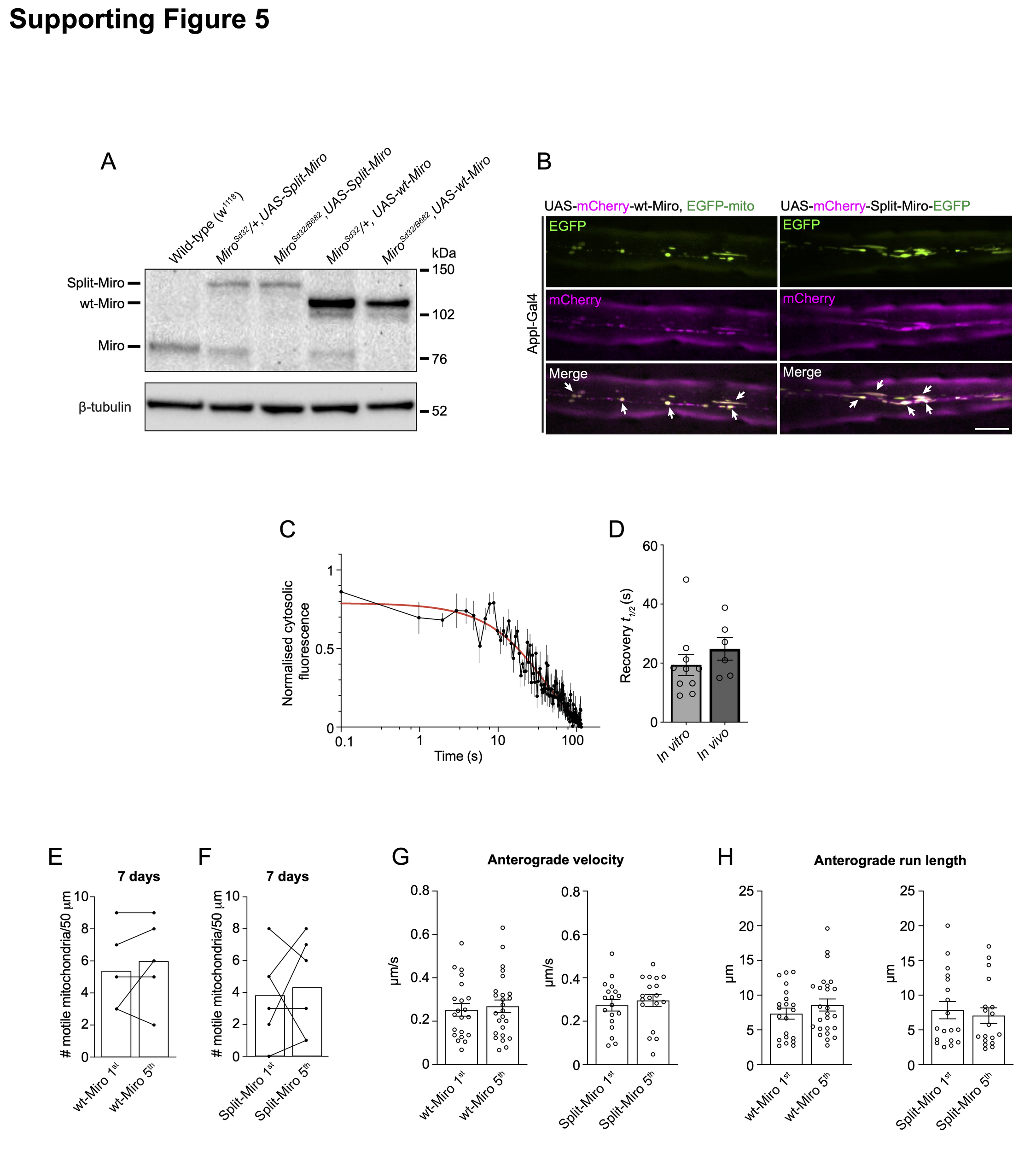

### Supporting Figure 6

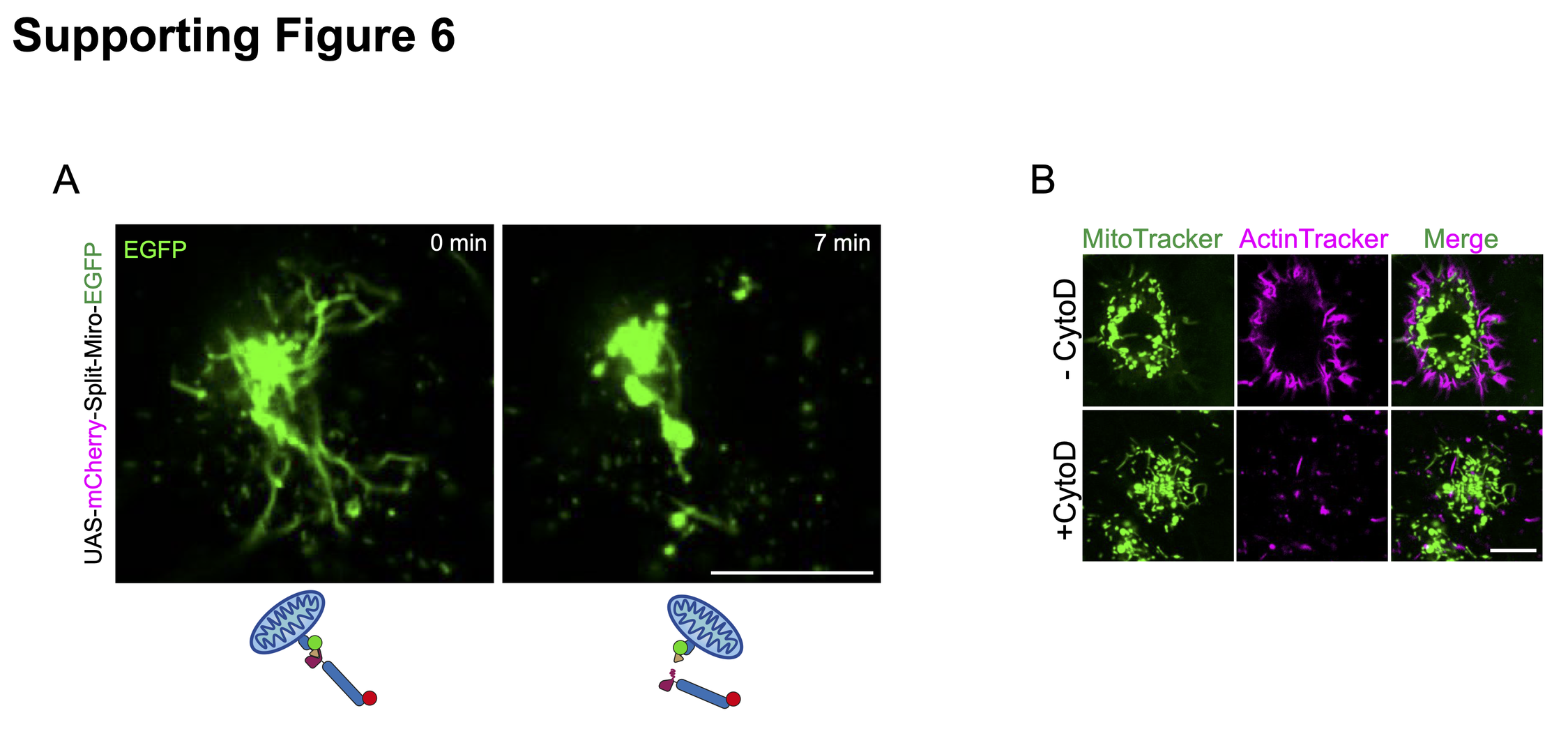
