## Supporting Tables 1-3 for "Optogenetic Miro cleavage reveals direct consequences of real-time loss of function in *Drosophila*"

**Supporting Table 1.** Plasmids used in this study.

| Construct number | Insert | Backbone | Cloning method, construct assembly and DNA sources |
|---|---|---|---|
| #1. | (Empty vector) | pAc5.1/V5-HisB (pAc5.1) | Gift of Alex Whitworth (MRC-MBU, Cambridge, UK). |
| #2. | (Empty vector) | pUASTattB | Gift of Manolis Fanto (King's College London, London, UK). |
| #3 | SPLICS-L | pT2-DsRed-UAS | Gift of Tito Calì (University of Padova, Padova, Italy). |
| #4 | $^{\text{KpnI}}$mCherry-Miro$^{\text{XbaI}}$ | pAc5.1/V5-HisB | NEBuilder HiFi DNA Assembly using primers #13,#14 (for mCherry) and #8,#2 (for Miro).<br>mCherry was PCR amplified from Addgene #81041 and fused to the *Drosophila* Miro-RE/RF cDNA sequence. |
| #5 | | pUASTattB | Restriction enzymes-based cloning. Insert cut and pasted from construct #4. |
| #6 | $^{\text{KpnI}}$EGFP-MiroN-LOV2 T406-407A$^{\text{NotI}}$ | pAc5.1/V5-HisB | NEBuilder HiFi DNA Assembly using primers #6,#7 (for EGFP); #8,#9 (for MiroN) and #10,#11 (for LOV2 T406-407A).<br>EGFP was PCR amplified from Clontech pEGFP-N1. MiroN, encompassing the aa 1-642 of *Drosophila* Miro, was PCR amplified from endogenous Miro- |

| | | | |
|---|---|---|---|
| | | | RE/RF. LOV2 T406-407A was obtained by site-directed mutagenesis of mCherry-MiroN-LOV2 wild type (wt). |
| #7 | $^{KpnI}$mCherry-MiroN-LOV2 T406-407A$^{NotI}$ | pAc5.1/V5-HisB | NEBuilder HiFi DNA Assembly using primers #13,#14 (for mCherry) and #8,#11 (for MiroN-LOV2 T406-407A).<br><br>mCherry was PCR amplified from Addgene #81057. MiroN-LOV2 T406-407A was amplified from EGFP-MiroN-LOV2 T406-407A (construct #6). |
| #8 | | pUASTattB | Restriction enzymes-based cloning. Insert cut and pasted from construct #7. |
| #9 | mCherry-MiroN-LOV2 wt | pAc5.1/V5-HisB | The construct was obtained by site-directed mutagenesis of construct #7 using primers #23, #24. |
| #10 | $^{NotI}$mCherry-Zdk1-MiroC$^{XbaI}$ | pAc5.1/V5-HisB | NEBuilder HiFi DNA Assembly using primers #3,#4 (for mCherry-Zdk1) and #5,#2 (for MiroC).<br><br>mCherry-Zdk1 was PCR amplified from Addgene #81057 and fused to MiroC (encompassing the aa 643-674 of *Drosophila* Miro). |

| | | | |
|---|---|---|---|
| #11 | $^{\text{KpnI}}$EGFP-Zdk1-MiroC$^{\text{XbaI}}$ | pAc5.1/V5-HisB | NEBuilder HiFi DNA Assembly using primers #6,#15 (for EGFP) and #16,#2 (for Zdk1-MiroC).<br>EGFP was PCR amplified from Clontech pEGFP-N1 and fused to Zdk1-MiroC amplified from construct #10. |
| #12 | | pUASTattB | Restriction enzymes-based cloning. Insert cut and pasted from construct #11. |
| #13 | $^{\text{NotI}}$Zdk1-MiroC$^{\text{XbaI}}$ | pAc5.1/V5-HisB | NEBuilder HiFi DNA Assembly using primers #12,#2.<br>Zdk1-MiroC was amplified from construct #10. |
| #14 | | pUASTattB | Restriction enzymes-based cloning. Insert cut and pasted from construct #13. |
| #15 | GFP-SKL | pGG101 | AcGFP-SKL (gift of Gohta Goshima, Nagoya University, Nagoya, Japan) |
| #16 | $^{\text{KpnI}}$Mito4xGCaMP6f$^{\text{NotI}}$ | pAc5.1/V5-HisB | NEBuilder HiFi DNA Assembly using primers #17,#18.<br>Insert PCR-amplified from Addgene #127870. |
| #17 | $^{\text{KpnI}}$EGFP$^{\text{EcoRI}}$SNPH$^{\text{XbaI}}$ | pAc5.1/V5-HisB | Restriction enzymes-based cloning. Human SNPH (Gift of Zu-Hang Sheng, NIH, USA) was cut |

| | | | |
|---|---|---|---|
| | | | and pasted into an EGFP-pAc5.1/V5-HisB vector. |
| #18 | $^{\text{KpnI}}$EBFP$^{\text{EcoRI}}$SNPH$^{\text{XbaI}}$ | pAc5.1/V5-HisB | NEBuilder HiFi DNA Assembly using primers #19,#20 for EBFP.<br>EBFP was PCR amplified from Clontech pEBFP-C1 (gift of Marc-David Ruepp, King’s College London, London, UK) and fused to human SNPH (Gift of Zu-Hang Sheng, NIH, Bethesda, USA). |
| #19 | $^{\text{BamHI}}$mito-ER.SPLICS$^{\text{XbaI}}$ | pAc5.1/V5-HisB | Restriction enzymes-based cloning. Mito-ER.SPLICS was cut and pasted from construct #3 into the pAc5.1/V5-HisB vector. |

**Supporting Table 2.**

| Primer number | Primer name | Direction | Sequence (5'-3') |
|---|---|---|---|
| NEBuilder HiFi DNA Assembly | | | |
| #1. | Miro (overlap to pAc5.1) | Fwd | **TCCAGAGACCCCGGATCGGG**GTACCATGGGACAGTACACGGCGTCGCAGCGCAAG |
| #2. | Miro (overlap to pAc5.1) | Rvs | **ACCTTCGAACCGCGGGCCCT**CTAGACTAACGGGTGTGGGCTCCGGCAGCACTTAT |
| #3. | mCherry (overlap to pAc5.1) | Fwd | **CAGATATCCAGCACAGTGGC**GGCCGCATGGTGAGCAAGGGCGAGGAGGATAACATG |
| #4. | Zdk1 (overlap to Miro-Cterm) | Rvs | **AGTCCCGCCTTTAACCACAG**GCTACCACCAGAACCACCTTTTGGGGCCTG |
| #5. | Miro-Cterm (overlap to Zdk1) | Fwd | **AAGGTGGTTCTGGTGGTAGC**CTGTGGTTAAAGGCGGGACTAGGAGTGGCC |
| #6. | EGFP (overlap to pAc5.1) | Fwd | **TCCAGAGACCCCGGATCGGG**GTACCATGGTGAGCAAGGGCGAGGAGCTGTTCACC |

| | | | |
|---|---|---|---|
| #7. | EGFP (overlap to Miro-Nterm) | Rvs | **GACGCCGTGTACTGTCCCAT**CTTGTACAGCTCGTCCATGCCGAGAGTGAT |
| #8. | MiroNterm (overlap to EGFP) | Fwd | **GCATGGACGAGCTGTACAAG**ATGGGACAGTACACGGCGTCGCAGCGCAAG |
| #9. | MiroNterm (overlap to LOV2) | Rvs | **GCAGCCAAGGATCCAGAACC**CTTGGGGTCCTCCGTCATCAGGCCGAATTG |
| #10. | LOV2 (overlap to Miro-Nterm) | Fwd | **TGATGACGGAGGACCCCAAG**GGTTCTGGATCCTTGGCTGCTGCACTTGAA |
| #11. | LOV2 (overlap to pAc5.1) | Rvs | **CGGGCCCTCTAGACTCGAGC**GGCCGCTTAAAGTTCTTTTGCCGCCTCATCAATATT |
| #12. | Zdk (overlap to pAc5.1) | Fwd | **CAGATATCCAGCACAGTGGC**GGCCGCATGGGTTCTGGATCCATGGTGGATAACAAATTC |
| #13. | mCherry_2 (overlap to pAc5.1) | Fwd | **TCCAGAGACCCCGGATCGGG**GTACCATGGTGAGCAAGGGCGAGGAGGATAACATG |

| | | | |
|---|---|---|---|
| #14. | mCherry (overlap to Miro-Nterm) | Rvs | **GACGCCGTGTACTGTCCCAT**CTTGTACAGCTCGTCCATGCCGCCGGTGGA |
| #15. | EGFP (overlap to Zdk1) | Rvs | **TCCACCATGGATCCAGAACC**CTTGTACAGCTCGTCCATGCCGAGAGTGAT |
| #16. | Zdk1 (overlap to EGFP) | Fwd | **GCATGGACGAGCTGTACAAG**GGTTCTGGATCCATGGTGGATAACAAATTC |
| #17. | Mito4xGCaMP6f (overlap to pAc5.1) | Fwd | **TCCAGAGACCCCGGATCGGG**GTACCATGAGCGTGCTGACACCTCTGCTGC |
| #18. | Mito4xGCaMP6f (overlap to pAc5.1) | Rvs | **CGGGCCCTCTAGACTCGAGC**GGCCGCTCACTTGGCGGTCATCATCTGGACAAACTC |
| #19. | EBFP (overlap to pAc5.1) | Fwd | **TCCAGAGACCCCGGATCGGG**GTACCATGGTGAGCAAGGGCGAGGAGCTGTTC |
| #20. | EBFP (overlap to SNPH) | Rvs | **GGCCGCTGCCCGGCATGGTG**AATTCCTTGTACAGCTCGTCCATGCCGAGAGTG |
| Site-directed mutagenesis | | | |

| | | | |
|---|---|---|---|
| #21. | LOV2 mut | Fwd | ATCCTTGGCT**G**CT**G**CACTTGAACGTA |
| #22. | LOV2 mut | Rvs | CCAGAACCCTTGTACAGC |
| #23. | LOV2 wt | Fwd | ATCCTTGGCT**A**CT**A**CACTTGAACGTA |
| #24. | LOV2 wt | Rvs | CCAGAACCCTTGGGGTCC |
| dsRNA template production | | | |
| #25. | Miro dsRNA | Fwd | TAATACGACTCACTATAGGGAGGGGAATTCACTAGGATAAGGGGA |
| #26. | Miro dsRNA | Rvs | ATTATGCTGAGTGATATCCCTCGCCATTAAATATCACTATATGTTAATCCA |
| #27. | Control dsRNA | Fwd | TAATACGACTCACTATAGGGAG |
| #28. | Control dsRNA | Rvs | ATTATGCTGAGTGATATCCCTCCTGTAGCCCAAGTTGTTGATATTAT |

**Supporting Table 3.**

| Strain | Source |
|---|---|
| Oregon-R | Bloomington Drosophila Stock Center (BDSC), #5 |
| $w^{1118}$ | BDSC, #6326 |
| UAS-mito::GFP | BDSC, #8442 |
| Appl-Gal4 | BDSC, #32040 |
| $miro^{B682}$/TM6c | BDSC, #52003 |
| $miro^{Sd32}$/T(2;3),CyO:TM6b | BDSC, #52002 |
| 20xUAS-GCaMP5G | BDSC, #56500 |
| w; 5xUAS-mCherry::MiroN-LOV2 T406-407A (attP40)/CyO | This study |
| w;; 5xUAS-EGFP::Zdk1-MiroC (attP2)/TM6c | This study |
| w;; 5xUAS-Zdk1-MiroC (attP2)/TM6c | This study |
| w; 5xUAS-Zdk1-MiroC (attP40)/CyO | This study |
| w; 5xUAS-mCherry::Miro wt (attP40)/CyO | This study |

| | |
|---|---|
| w;; 5xUAS-mCherry::Miro wt (attP2)/TM6c | This study |
| Genotypes for behavioural assay | |
| w, Appl-Gal4/Y; 5xUAS-mCherry::MiroN-LOV2 T406-407A (attP40)/+; 5xUAS-Zdk1-MiroC (attP2), miro$^{Sd32}$/miro$^{B682}$ | |
| w, Appl-Gal4/Y; +/+; 5xUAS-mCherry::Miro wt (attP2), miro$^{Sd32}$/miro$^{B682}$ | |
| w, Appl-Gal4/Y; 5xUAS-mCherry::MiroN-LOV2 T406-407A (attP40)/+; 5xUAS-Zdk1-MiroC (attP2), miro$^{Sd32}$/EGFP-SNPH (attP2), miro$^{B682}$ | |
| Genotypes for live imaging of mitochondrial transport | |
| w, Appl-Gal4/+; UAS-mito::GFP/+; 5xUAS-mCherry::Miro wt (attP2), miro$^{Sd32}$/+ | |
| w, Appl-Gal4/+; 5xUAS-mCherry::MiroN-LOV2 T406-407A (attP40)/ +; 5xUAS-EGFP-Zdk1-MiroC (attP2), miro$^{Sd32}$/+ | |
| Genotypes for live imaging of neuronal activity | |
| w, Appl-Gal4/+; 5xUAS-mCherry::Miro wt (attP40)/+; 5xUAS-Zdk1-MiroC (attP2), miro$^{Sd32}$/20xUAS-GCaMP5G | |
| w, Appl-Gal4/+; 5xUAS-mCherry::MiroN-LOV2 T406-407A (attP40)/+; 5xUAS-Zdk1-MiroC (attP2), miro$^{Sd32}$/20xUAS-GCaMP5G | |
